# Multiplexed sgRNA Expression Allows Versatile Single Non-repetitive DNA Labeling and Endogenous Gene Regulation

**DOI:** 10.1101/121905

**Authors:** Shipeng Shao, Lei Chang, Yuao Sun, Yingping Hou, Xiaoying Fan, Yujie Sun

## Abstract

The CRISPR/Cas9 system has made significant contribution to genome editing, gene regulation and chromatin studies in recent years. High-throughput and systematic investigations into the multiplexed biological systems and disease conditions require simultaneous expression and coordinated functioning of multiple sgRNAs. However, current co-transfection based sgRNA co-expression systems remain poorly efficient and virus-based transfection approaches are relatively costly and labor intensive. Here we established a vector-independent method allowing multiple sgRNA expression cassettes to be assembled in series into a single plasmid. This synthetic biology-based strategy excels in its efficiency, controllability and scalability. Taking the flexibility advantage of this all-in-one sgRNA expressing system, we further explored its applications in single non-repetitive genomic locus imaging as well as coordinated gene regulation in live cells. With its strong potency, our method will greatly facilitate the understandings in genome structure, function and dynamics, and will contribute to the systemic investigations into complex physiological and pathological conditions.

## INTRODUCTION

Since its discovery, the clustered regularly inter-spaced short palindromic repeats (CRISPR)-associated Cas9 nuclease has largely revolutionized the fields in genome editing, gene regulation and chromatin studies in live cells (1). Compared to protein-based recognition systems such as ZFN and TALE, CRISPR recognizes target DNA via Watson-Crick base pairing, allowing it to perform high-throughput chromatin targeting in an effective and efficient way (2). Emerging new versions of the CRISPR/Cas9 system have been adopted to expand our capacity for precise genome manipulation (3-7). These powerful toolkits carry tremendous potentials in both biomedical researches and therapeutic genomic targeting (8).

Simultaneous expression and coordinated functioning of multiple sgRNAs in live cells are often necessary in circumstances such as simultaneous knockout (9) and regulation of multiple endogenous genes (10). Notably, CRISPR-based labeling and imaging of genomic loci with non-repetitive sequences in live cells also require to tile the loci with multiple sgRNAs (11).

There are several approaches available for simultaneous delivery of multiple sgRNAs to scale up the targeting capacity of the CRISPR/Cas9 system. For instance, cells can be transfected with a mixture of multiple sgRNA plasmids by chemi-transfection or electro-transfection (12,13). However, such transient cotransfection of different plasmids is relatively inefficient and also results in variable expression levels of each sgRNA in single cells. Similarly, delivery of multiple sgRNAs using lentivirus infection (11) can also lead to variable efficacy in delivery and expression. Alternatively, although microinjection can deliver multiple sgRNAs into a single cell without the limitation of plasmid type and number (14), it suffers from low throughput and requires special equipments and superb skills. With the evolution of synthetic biology, new cloning methods have emerged in the last two years to achieve high speed, efficiency and flexibility in assembly of multiple DNA fragments (15-17). For instance, the Golden Gate cloning method was used for fast assembly of TALE repeats (18) and for assembly of short serial sgRNAs (9,19). However, the number of sgRNA cassettes assembled in one reaction is quite limited following current protocols as the assembly efficiency decreases dramatically as the number of cassettes increases. Even though a recently developed hierarchical assembly method could assemble multiple sgRNAs, as it relies on receptor plasmids for every step, the whole procedure requires several rounds of transformation, verification, plasmid extraction, and thus on average takes more than 20 days to assembly 20 sgRNA expression cassettes into a single plasmid (20). Collectively, a well-rounded multiplexing sgRNA expression system needs to balance in ease of implementation, delivery efficiency, precise stoichiometry control, and cost of time.

In this work, we have developed a hierarchical cloning strategy to assemble multiple sgRNA expression cassettes into a single vector. We demonstrated that this new approach is not only robust and time-saving in assembly of different numbers of sgRNA expression cassettes at will but also able to assure precise control of stoichiometry of different sgRNAs. Using this all-in-one sgRNA expressing system, we successfully labeled and imaged non-repetitive genomic loci, MUC4 and HER2 genes, in living human cells. Additionally, we demonstrated that the all-in-one sgRNA expressing system is highly efficient in gene activation and repression. Using multiple sgRNAs to tile the promoter of endogenous gene SOX2 to recruit multiple transcriptional regulators, we found that the regulation efficiency by this synergistic gene regulation approach was superior to that by single sgRNA and co-transfection of multiple sgRNAs. We also showed that the all-in-one sgRNA expressing system allows parallel expression of different sgRNAs with relatively stable stoichiometry in their expression level, leading to reliable simultaneous activation and/or repression of multiple endogenous genes in the same cell. In summary, our modular sgRNA assembly system can be readily automated and will be highly useful for applications in cell biology and genetics, such as gene editing, gene regulation, pathway engineering and live cell chromatin labeling.

## MATERIALS & METHODS

### Hierarchical assembly of multiple sgRNA expression cassettes

All primers were ordered from Ruibo Biotech (Beijing). The U6 promoter, conventional and modified sgRNA expression cassettes were amplified from plasmids. The oligoes (F + R, 100 μΜ each) for each targeting sgRNA were phosphorylated by T4 PNK (NEB) and annealed in a thermocycler by using the following parameters: 37 °C for 30 min; 95 °C for 5 min; ramp down to 25 °C at 5 °C min^-1^. The phosphorylated and annealed oligoes were then diluted to 1:100 by adding 1 μl oligoes to 99 μl ddH_2_O.

For the first assembly step, U6 promoter (10 ng), annealed oligoes (1 μΜ) and sgRNA scaffold (5 ng) were mixed with BsmBI (1 μl, NEB) and T4 ligase (0.5 μl, NEB) together for Golden Gate reaction in 20 separate tubes by using the following parameters: 37 °C for 10 min; 16 °C for 10 min; 10 cycle; 55 °C for 10 min; 85 °C for 10 min. The reaction mix was then diluted by 200 folds in ddH_2_O. 1 μl diluted mix was used as template for PCR reaction. The primer sequence and allocation have been described in Supplementary Tables 1 and 2. After the PCR reaction, gel electrophoresis was used to check the reaction. After gel extraction, a QIAquick gel extraction kit was used to ensure that the product contains only the intended U6+oligos+sgRNA ligation product.

For the second assembly step, the 20 ligation fragments (10 ng each) were mixed in 4 separate tubes (1-5 in tube #1, 6-10 in tube #2, 11-15 in tube #3, 16-20 in tube #4) with BsaI (1 μl, NEB) and T4 ligase (0.5 μl, NEB) together for Golden Gate reaction by using the following parameters: 37 °C for 10 min; 16 °C for 10 min; 10 cycle; 37 °C for 10 min; 85 °C for 10 min. The reaction mix was then diluted by 200 folds in ddH_2_O. 1 μl diluted mix was used as template for the PCR reaction. The primer sequences and allocation were described in Supplementary Tables 1 and 2. After the PCR reaction, gel electrophoresis was used to check the reaction. After gel extraction, a QIAquick gel extraction kit was used to ensure that the product contains only the intended 5 ligation length product. This PCR step needs to be optimized to ensure the maximum productivity due to the repetitive sequence of the 5 sgRNA expression cassettes. If the intermediates (5 sgRNA expression plasmids) were needed, the Golden Gate reaction should be performed with the corresponding receptor plasmids.

For the third assembly step, the 4 ligated fragments (100 ng each) and 50 ng receptor plasmid pMulti-sgRNA-LacZα (home-made, Supplementary Map 1 & 2) were mixed with BpiI (1 μl, Thermo Fisher Scientific) and T4 ligase (0.5 μl, NEB) together for Golden Gate reaction by using the following parameters: 37 °C for 10 min; 16 °C for 10 min; 10 cycle; 37 °C for 10 min; 85 °C for 10 min. The competent *E. coli* strain *Stbl3* (TransGen Biotech) was transformed with the final reaction mix and cultured in 30 °C to minimize the opportunity of losing the repeat units. The plasmids were linearized and checked by gel electrophoresis to ensure that the final plasmid contains the proper length of repeats.

### Construction of other plasmids used in this work

The NLS_Sv40_-dCas9-(NLS_Sv40_)_x3_-(GCN4_-v4_)_x24_-NLS_Sv40_-P2A-BFP fragment was amplified by PCR from plasmid pHRdSV40-NLS-dCas9-(GCN4_-V4_)_X24_-NLS-P2A-BFP-dWPRE (Addgene Plasmid #60910) and then ligated into PiggyBac plasmid pB-TRE3G-BsmBI-EF1α-PuroR-P2A-rtTA (home-made) by Golden Gate assembly with BsmBI and T4 ligase (NEB). The scFV-sfGFP-GB1-NLS_SV40_ fragment was amplified from plasmid pHR-scFv-GCN4-sfGFP-GB1-NLS-dWPRE (Addgene Plasmid #60906) and then ligated into PiggyBac plasmid pB-TRE3G-BsmBI-EF1α-HygroR-P2A-rtTA (home-made) by Golden Gate assembly with BsmBI and T4 ligase (NEB). The tdMCP fragment was amplified from phage-ubc-nls-ha-tdMCP-gfp (Addgene Plasmid #40649), and the KRAB was amplified from pHAGE-EF1α-dCas9-KRAB (Addgene Plasmid #40649) and then ligated into plasmid pB-CMV (home-made). The tdPCP fragment was amplified from phage-ubc-nls-ha-tdPCP-gfp (Addgene Plasmid #40650), and the VPR was amplified from SP-dCas9-VPR (Addgene Plasmid #63798) and then ligated into plasmid pB-CMV (home-made). The telomere, rDNA10, MUC4-E2 targeting sgRNAs and other sgRNA expression plasmids were made according to procedures described previously (6). All primers used in this work were listed in Supplementary Tables 1-8.

### Cell Culture, Transfection, Stable Cell Line Construction

Human breast cancer cell line MDA-MB-231 was maintained in Dulbecco’s modified Eagle medium (DMEM) with high glucose (Lifetech), 10% FBS (Lifetech), 1X penicillin/streptomycin (Lifetech). All cells were maintained at 37 °C and 5% CO_2_ in a humidified incubator. All plasmids were transient transfected with Lipofectamine 2000 (Lifetech) in accordance with the manufacturer’s protocol. To construct a stable cell line, MDA-MB-231 cells were spread onto a 6-well plate one day before transfection. On the next day, the cells were transfected with 500 ng pB-dCas9-(GCN4)_X24,_ 500 ng pB-scFV-sfGFP, and 200 ng pCAG-hyPBase using Lipo 2000. Hygromycin (200 μg/ml) and puromycin (5 μg/ml) were used for the selection for about 2 weeks. Cells with proper expression level of Cas9-(GCN4)_X24_ and scFV-sfGFP were selected using FACS.

### FISH and Immunofluorescence

The transfected cells were washed with PBS briefly once and incubated with extraction buffer (0.1 M PIPS, 1 mM EGTA, 1 mM MgCl_2_, 0.2% TritonX-100) for 1 min to extract cytoplasm diffusive proteins. Cells were then washed once with PBS and fixed with 4% PFA for 15 minutes. Cells were permeabilized with 0.1% saponin/0.1% Triton X-100 in PBS for 10 min at RT (room temperature) and then washed twice in PBS for 5 min at RT. Cells were incubated for 1 h in 20% glycerol/ PBS at RT and then frozen/ thawed 3 times in liquid nitrogen. Cells were washed twice in PBS for 5 min at RT, incubated in 0.1 M HCl for 30 min at RT and then washed in PBS for 5 min at RT. Cells were permeabilized with 0.5% saponin/0.5% Triton X-100 in PBS for 30 min at RT and washed twice in PBS for 5 min at RT. Cells were equilibrated in 50% formamide/2x SSC for 20 min at RT. The probe was added in hybridization buffer and the dish was warmed to 78 °C for precisely 2 min. Cells were incubated overnight at 37 °C in a dark humidified chamber. The next day, cells were washed in the following buffer sequentially, 50% formamide/2X SSC for 15 min at 45 °C, 0.2X SSC for 15 min at 63 °C, 2X SSC for 5 min at 45 °C. and 2X SSC for 5 min at RT.

For immunofluorescence, cells were incubated in 0.5% Triton followed by 30 min 5% BSA blocking at RT. The cells were then stained with human anti-GFP antibodies (proteintech #66002-1-Ig) in blocking buffer for 60 min, washed with PBS, and then stained with Cy3-labeled secondary antibodies in blocking buffer for 1 h. The labeled cells were washed with PBS, post-fixed with 4% PFA for 10 min at RT. Lastly, the cells were stained with DAPI (5 μg/ml in PBS) for 2 min in at RT and then wash 3 times with PBS.

### qRT–PCR

About 48h post transfection in six-well plates with proper amount of plasmids, total RNA were extracted by TRIzol (Invitrogen). Complementary DNAs (cDNAs) were synthesized using RevertAid First Strand cDNA synthesis Kit (Thermo Fisher Scientific) and qPCR was performed using SYBR Green PCR Master Mix (Roche). GADPH was used as the internal reference. All primers used in the qPCR step were designed by primer-BLSAT on NCBI website. For sequences of the primers, see Supplementary Table 7.

### Fluorescence-activated cell sorting (FACS) analysis

Cells were trypsinized 48 h after transfection and centrifuged at 300xg for 5 min at RT. The supernatant was removed, and the cells were resuspended in 1× PBS. Fortessa flow analyzers (BD Biosciences) were used for fluorescence-activated cell sorting analysis with the following settings. BFP was measured using a 405 nm laser and a 450/50 filter and sfGFP was measured using a 488 nm laser and a 530/30 filter. For each sample, about 10^4^ cell events were collected. The FACS results were analyzed by FlowJo.

### Image acquisition and analysis

Briefly, all images were taken on a Nikon Ti-E inverted microscope equipped with a 100X UPlanSApo, N.A. = 1.40, oil-immersion phase objective and EMCCD. The microscope stage incubation chamber was maintained at 37 °C and 5% CO_2_. A 405-nm laser was used to excite BFP and DAPI; a 488-nm laser was used to excite sfGFP; a 561-nm laser was used to excite Cy3; a 647-nm laser was used to excite Cy5. The laser power was modulated by an acousto-optic tunable filter (AOTF). Movies of short time scale chromatin loci dynamics in living cells were acquired at 20 Hz.

Analysis of MSD and velocity autocorrelation was carried out using custom MATLAB scripts. The MSD analysis method has been described previously (6). Average velocity autocorrelation function was calculated as described previously by Joseph S. Lucas et al. (21). The function can be expressed as follows:

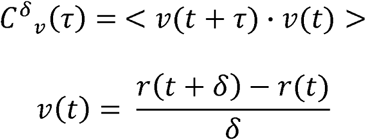
where δ is the time interval over which the average velocity is calculated and τ is the lag time over which the correlation in velocities is calculated.

## RESULTS

### Plasmid-independent hierarchical assembly of multiple sgRNA expression cassettes

To achieve high-throughput assembly of multiple sgRNA cassettes, here we have developed a plasmid-independent, PCR-based hierarchical assembly strategy. Using a series of three one-pot Golden Gate cloning with type II restriction endonuclease, we proved the robustness and high efficiency of our method in assembly of repeated units (Figure 1A). Firstly, PCR-amplified conventional or functionally modified sgRNA scaffolds and U6 promoter were ligated with respectively synthesized targeting oligoes by Golden Gate cloning (Supplementary Table 9). Then the reaction mix was amplified by PCR with U6-forward and sgRNA-scaffold-reverse primers without purification (Figure 1B, Supplementary Table 1). Thirdly, we designed junctions allowing ordered ligation to form the intermediate fragments. We used these junctions to assemble five sgRNA cassettes into one by a second round of Golden Gate cloning (Figure 1C, Supplementary Table 2). Lastly, a third Golden Gate reaction was performed to assemble the five-cassette-intermediates into the final construct carrying all 20 sgRNAs. The whole assembly procedure takes about 4-5 days. Random selection of end clones were verified (Figure 1D). Aside from high-efficiency, highspeed and high-throughput in generating a multiple-sgRNA cluster, our approach also provides the flexibility for assembling different numbers of sgRNAs using varied combinations of the forward and reverse primers.

**Figure 1.**
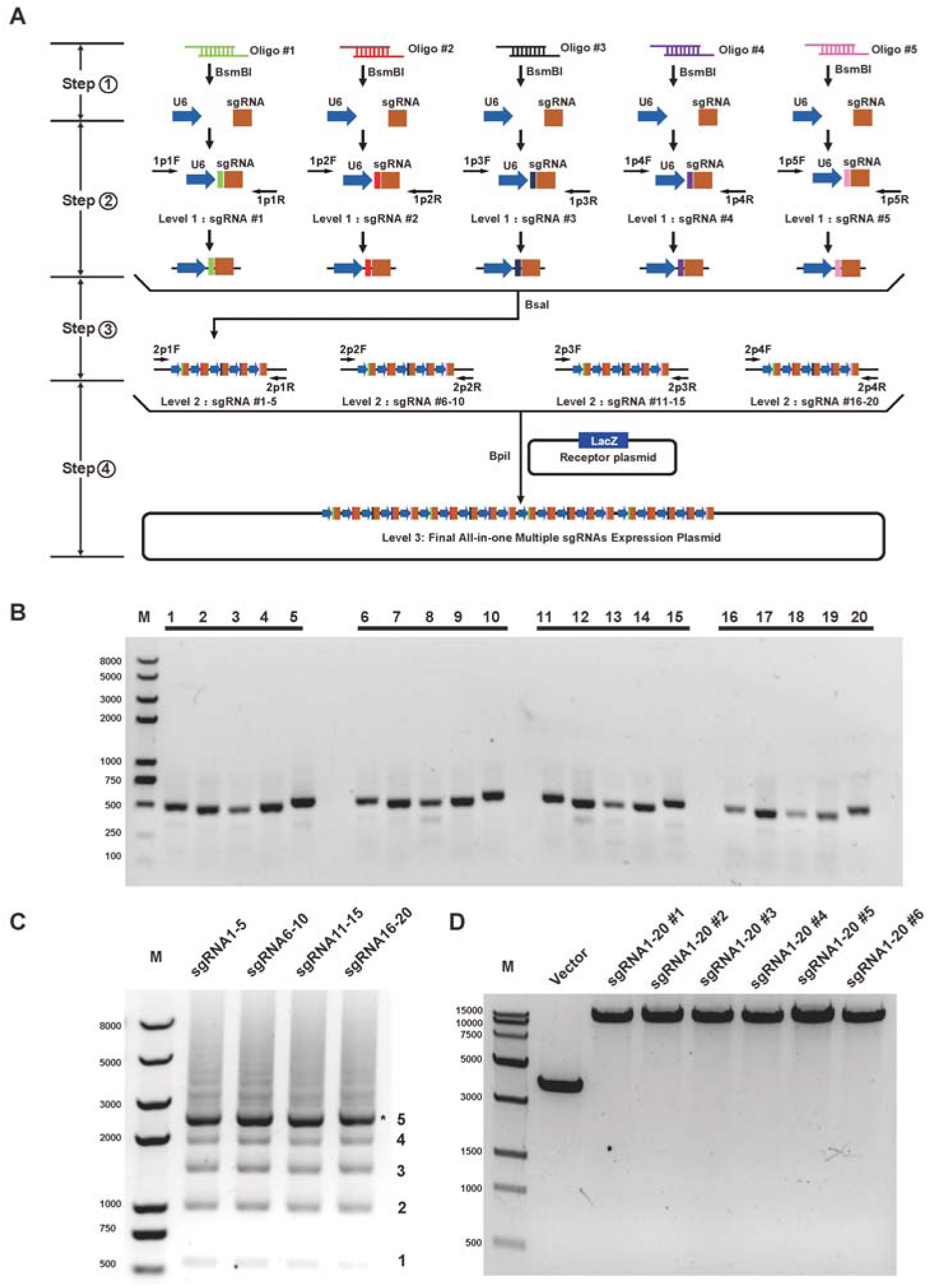
Hierarchical assembly of multiplexed sgRNA expression plasmid. (**A**). Workflow for the hierarchical assembly of 20 sgRNAs. U6 promoter and sgRNA scaffold are PCR amplified and then ligated with each annealed oligo in 20 separated Golden Gate reaction with a type IIs restriction endonuclease BsmBI (Step 1). Each Golden Gate reaction mix was then used as template for amplification without purification by 20 pair of primers, which can be grouped into 4 categories, each member in a group with different adaptors (Step 2). The purified 20 sgRNAs expression cassettes were then separated into 4 group and implemented with the second round Golden Gate reaction using BsaI (Step 3). The four fragments each carried 5 sgRNAs expression cassettes were purified and then ligated together into a third round Golden Gate reaction with a receptor plasmid using BpiI (Step 4). Due to the repetitive sequence of each sgRNA expression cassette, each step should be verified by agar gel electrophoresis to guarantee that each reaction was carried out successfully. (**B**). A typical gel of the first round PCR result for the successful ligation of U6, annealed oligoes, sgRNA scaffold. Trans2K plus II DNA ladder (TransGen Biotech) was shown in the left. The 20 individual sgRNA expression cascade all show the expected length on the gel, indicating that this cloning method is quite robust. A detail to note is that the first and fifth sgRNA in each group are slightly longer than the remaining one. This is because two handles are added to the 5’ of the first sgRNA and 3’ of the fifth sgRNA to facilitate the PCR of the five sgRNAs in the following step. (**C**). A typical gel of the second round PCR result for the successful ligation of five sgRNA expression cassettes into one fragment. Trans2K plus II DNA ladder (TransGen Biotech) was shown in the left. Due to the repetitive nature of the five sgRNAs fragment, laddering effect was observed. The indicated repeat number was shown in the right. (**D**). A typical gel of the verification of the final all-in-one plasmid. Trans15K DNA ladder (TransGen Biotech) was shown in the left. All the plasmids were linearized by endonuclease enzyme. The empty vector was also shown as a control. All the tested six plasmids have the right insertion size, suggesting the high efficiency of the assembly process.

### Establishment of a stable cell line expressing the SunTag system

We fused the SunTag signal amplification system (22) to dCas9 for the enhancement of fluorescence signal that one sgRNA can recruit. A previous study has shown that telomeres labeled with dCas9-SunTag were nearly 20-fold brighter compared to those labeled by dCas9 without alteration of the telomere mobility (22). However, labeling of non-repetitive chromosomal loci using dCas9-SunTag has not been realized. We reasoned that with dCas9-SunTag, simultaneous expression of multiple sgRNAs in a single cell can realize non-repetitive chromosomal locus labeling with high quality and efficiency (Figure 2A).

**Figure 2.**
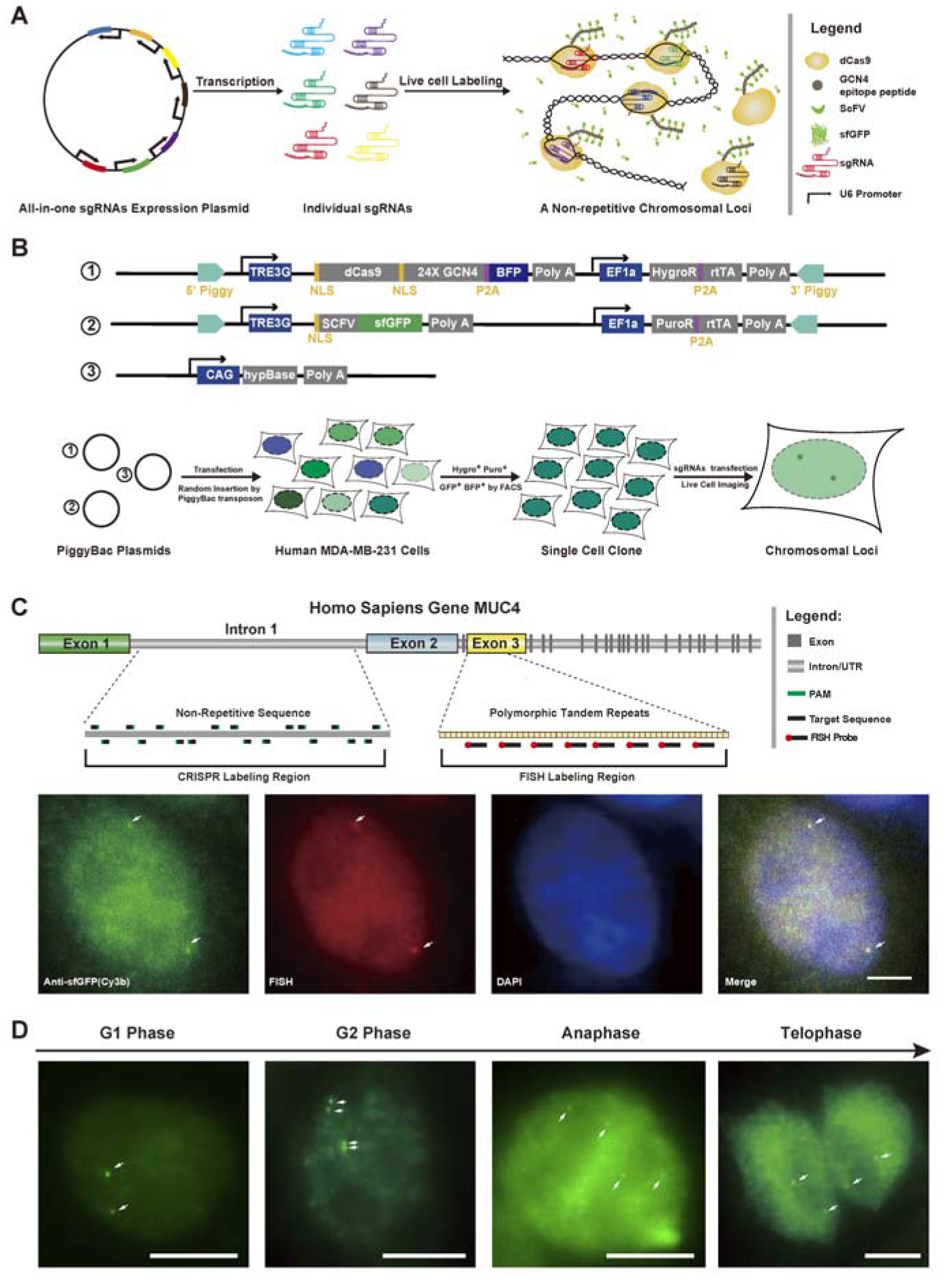
Single non-repetitive sequence labeling by multiplexed sgRNAs expression. (**A**). Schematic diagram of a non-repetitive sequence labeling of the genome. With the cell line that expressed proper level of SunTag system and 20 different sgRNAs tiling a short range sequence, arbitrary target loci can be labeled. The all-in-one plasmid carries 20 sgRNAs expression cassette each with a U6 promoter and poly U terminator so that individual sgRNA can be expressed independently. Six sgRNAs are shown as a demo here. The background fluorescent signal arises from the unbound dCas9 with multiple sfGFP molecules and the free sfGFP single molecules. Minimal level and proper stoichiometry (1:24) of SunTag system is the key point for the successful labeling. In principle, the 20 sgRNAs can direct 20 dCas9-sunTag molecule, that is 480 sfGFP molecules, to the target loci to gather the fluorescent signal. (**B**). Workflow for the establishment of live imaging system with proper expression level. Piggybac plasmids dCas9-(GCN4)_X24_-P2A-BFP and scFV-sfGFP were used to construct the stable cell line. The MDA-MB-231 cells were co-transfected with SunTag expression plasmids in PiggyBac backbone and transposase expression plasmid, followed by puromycin and hygromycin selection for about 2 weeks. sfGFP and BFP double positive single clones with minimal background expression and high labeling efficiency were selected and transient transfected with multiple sgRNA expression plasmid to test the performance of each clone in subsequent live imaging experiment. (**C**). The labeling result of non-repetitive sequence in human cells. By transfection of 20 sgRNAs from an all-in-one plasmid targeting the first intron of human MUC4 gene, bright fluorescent puncta (indicated by arrow) can be visualized. Fluorescence in situ hybridization against the third exon of MUC4 was used to verified the labeling result of CRISPR. The nucleus was stained with DAPI. The merged image showed the colonialization of CRISPR signal and FISH signal. All scale bars are 5 μm. (**D**). Single non-repetitive sequence can be labeled in different cell cycle stages. In G1 of interphase, two loci can be visualized. In G2 of interphase, four loci appeared as closely located pairs. During mitosis, the duplicated loci were allocated into two cells. In anaphase and telophase, the position of MUC4 loci nearly mirrored each other All scale bars are 10 μm.

Transient expression of two components of the SunTag system usually results in either incomplete labeling or excessive background from unbound molecules. In order to minimize those effects, we integrated the SunTag expression cassette into the genome using the PiggyBac transposon system (23) and generated a stable cell line with minimal expression level of dCas9-(GCN4)_X24-_P2A-BFP and scFV-sfGFP-GB1 (Figure 2B and Supplementary Figures 1& 2). With the low expression level, diffusing dCas9 single molecules were able to be observed in the GFP channel (Supplementary Figure 3A) and repetitive chromosomal loci, such as telomeres or the 2rd exon of MUC4 gene, were successfully labeled with dCas9-SunTag in live MDA-MB-231 cells (Supplementary Figure 3B & C). In order to compare the signal-to-noise ratio (SNR) between dCas9-GFP and the dCas9-SunTag system, the tandem array of 5S ribosomal DNA (5S rDNA) on chromosome 1, which consists of 17 repeats, was labeled and quantified (Supplementary Figure 4). The results showed that the SNR was improved nearly 6 folds with dCas9-SunTag. The significant improvement in SNR allows imaging with low laser power and short exposure time, enabling long-term tracking of chromosomal loci.

**Figure 3.**
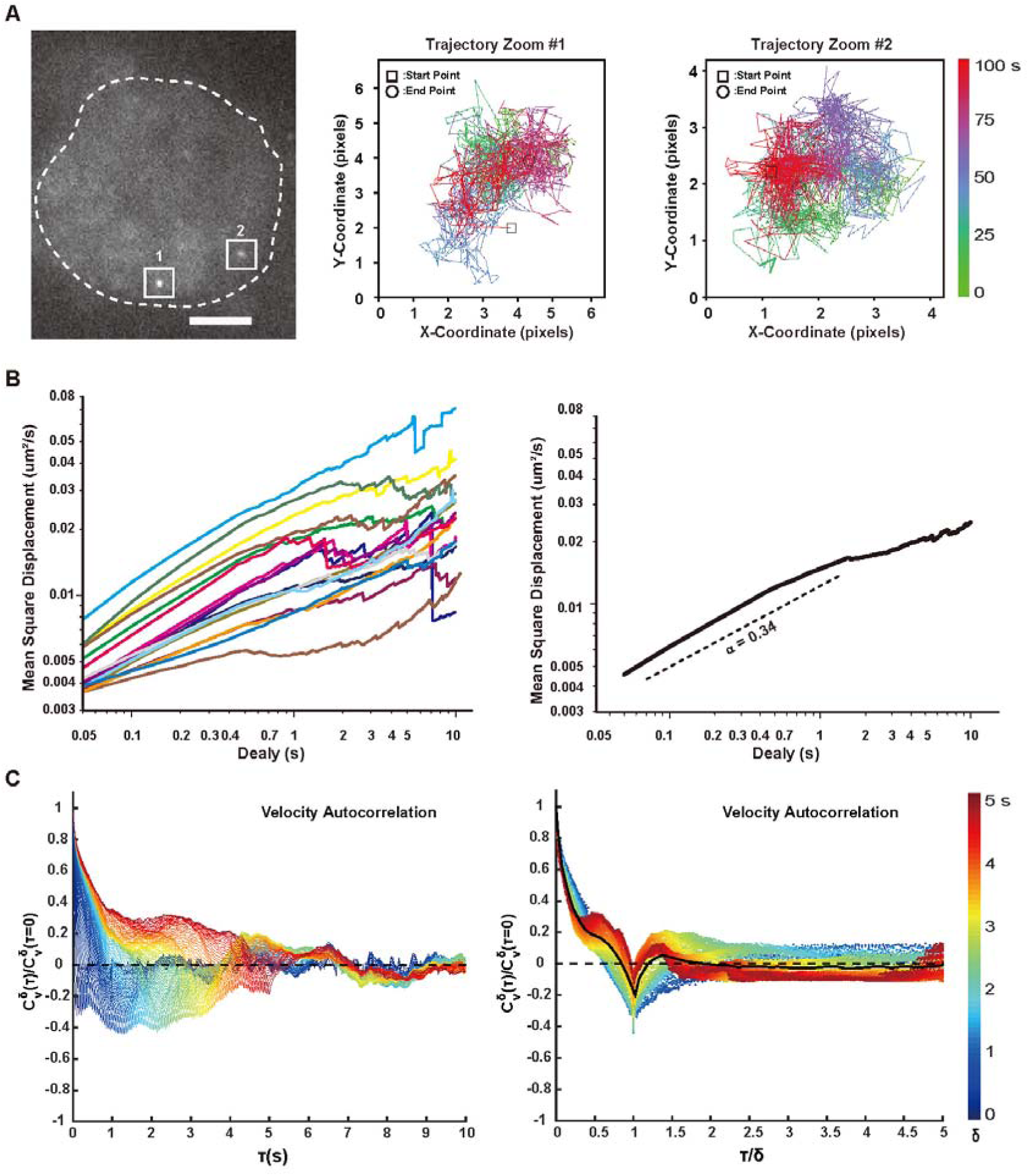
Live cell tracking of single non-repetitive genomic locus. (**A**). Typical image and trajectories of the MUC4 locus in live cell. Trajectories were displayed by the color-coded time points with green for the starting, blue for the intermediate and red for the ending frames. The time lookup table was in the right. Coordinate was measured by pixel. 1 pixel = 160 nm. Scale bar: 5 μm. (**B**). Time-averaged MSD plotted as a function of time lag (delay) for the MUC4 region in live cells. MSD was shown for delay up to 10 % of total imaging time. 17 trajectories were analyzed individually (left) or averaged among all the trajectories (right). Dashed lines indicate a subdiffusive scaling exponent (α) of 0.34, demonstrating the locus was undergoing constraint diffusion. (**C**). Velocity autocorrelation analysis of MUC4 locus in live cells. Velocity was calculated over discretization intervals (δ) ranging from 0.05 s to 5 s in 0.05 s intervals (left). Velocity autocorrelation curves for different values of δ, plotted against a rescaled timelag (t/δ) (right). The color scheme represents the values of δ from small (blue) to large (red).

### Single non-repetitive sequence labeling by multiplexed sgRNA expression

We next tried the multiple sgRNA expressing strategy for non-repetitive sequence labeling in living cells. We first chose MUC4 gene as a demonstration as it has tandem repeat regions advantageous for the validation by FISH. Specifically, 20 sgRNAs were deigned against a stretch of DNA (about 1.5 kb) and cloned into one plasmid by the hierarchical assembly method (Supplementary Table 3). The SunTag cell line was transfected with the all-in-one plasmid and imaged 24 h after transfection.

To verify the specificity and efficiency of non-repetitive MUC4 gene labeling, we performed two-color imaging with CRISPR probes targeting the non-repetitive sequence in the first intron of MUC4 and FISH probes targeting the polymorphic tandem repeats in the third exon. Because the harsh conditions of FISH sample preparation disrupted the fluorescence signal of GFP, we performed immunofluorescence against GFP. Colocolization of the two channels proved that the CRISPR probes indeed labeled the MUC4 locus (Figure 2C).

By labeling specific non-repetitive genomic loci with CRISPR, we could investigate chromosome reorganization at different cell cycle stages (Figure 2D & Supplementary Figure 5). Similar results were also obtained for another non-repetitive gene HER2 (Supplementary Figure 6). These results indicate that combination of the all-in-one sgRNA expression plasmid and SunTag signal amplification system can be used to label arbitrary sequences in the genome, visualize their positions in the nucleus, and study the relationship between the gene position and its expression (24).

We further tracked the dynamics of the MUC4 locus labeled with dCas9-SugTag (Supplementary Movie 1). Two typical trajectories in the same cell were observed (Figure 3A). No obvious correlation was found between the two allelic loci, indicating the heterogeneity of the surroundings of allelic genes. We calculated the average mean square displacement of 17 different loci and extracted the anomalous diffusion scaling exponents (α = 0.34) (Figure 3B), indicating that the movement of chromosomal loci is subdiffusive.

To obtain a more robust measurement of the subdiffusive behavior of the MUC4 locus, we calculated the average velocity autocorrelation function *C* ^δ^_V(T)._ This function describes what degree the average velocity over a time interval δ is correlated with the average velocity over another time interval δ that is separated by τ from the first one (21). The correlation function *C* ^δ^_V(T)_ was calculated for different values of δ and plotted as a function of the time lag τ. For all δ, *C* ^δ^_V(T)_ dropped into negative values before finally decaying to zero, indicating the negative correlation of MUC4 loci (Figure 3C, left). We then analyzed the behavior of *C* ^δ^_V(T)_ as a function of the ratio (τ/δ), and a single master curve was shown (Figure 3C, right). This indicates that the motion of MUC4 segments possessed the same negative correlation property at different temporal scales. Taken together, the results indicate that the MUC4 locus underwent a ‘back and forward’ movement, *i.e*. motion in one direction is likely to be followed by motion in the opposite direction, which may arise from restoring forces of its surrounding chromatin fiber.

### Synergistic regulation of endogenous genes by multiplexed sgRNA expression

Besides the single non-repetitive DNA sequence imaging, the multiplexed sgRNA expression strategy can also be used for synergistic regulation of endogenous gens as multiple sgRNAs can recruit multiple copies of proteins to function synergistically. We designed sgRNAs targeting the human SOX2 gene. Both the assembly intermediate (5 sgRNAs) and final product (20 sgRNAs) were inspected (Figure 4A & C). We found that the efficiency of synergistic activation of endogenous genes was much higher than that of single sgRNA-mediated activation or cotransfection of multiple sgRNAs. Furthermore, similar results were obtained for Cas9-KRAB mediated SOX2 repression experiments (Figure 4B & D). It needs to note that the activation or repression effect was not in proportion to the number of sgRNA. In addition, binding of multiple sgRNAs to the promoter region of a transcribed gene may also interfere binding of other transcription factors.

**Figure 4.**
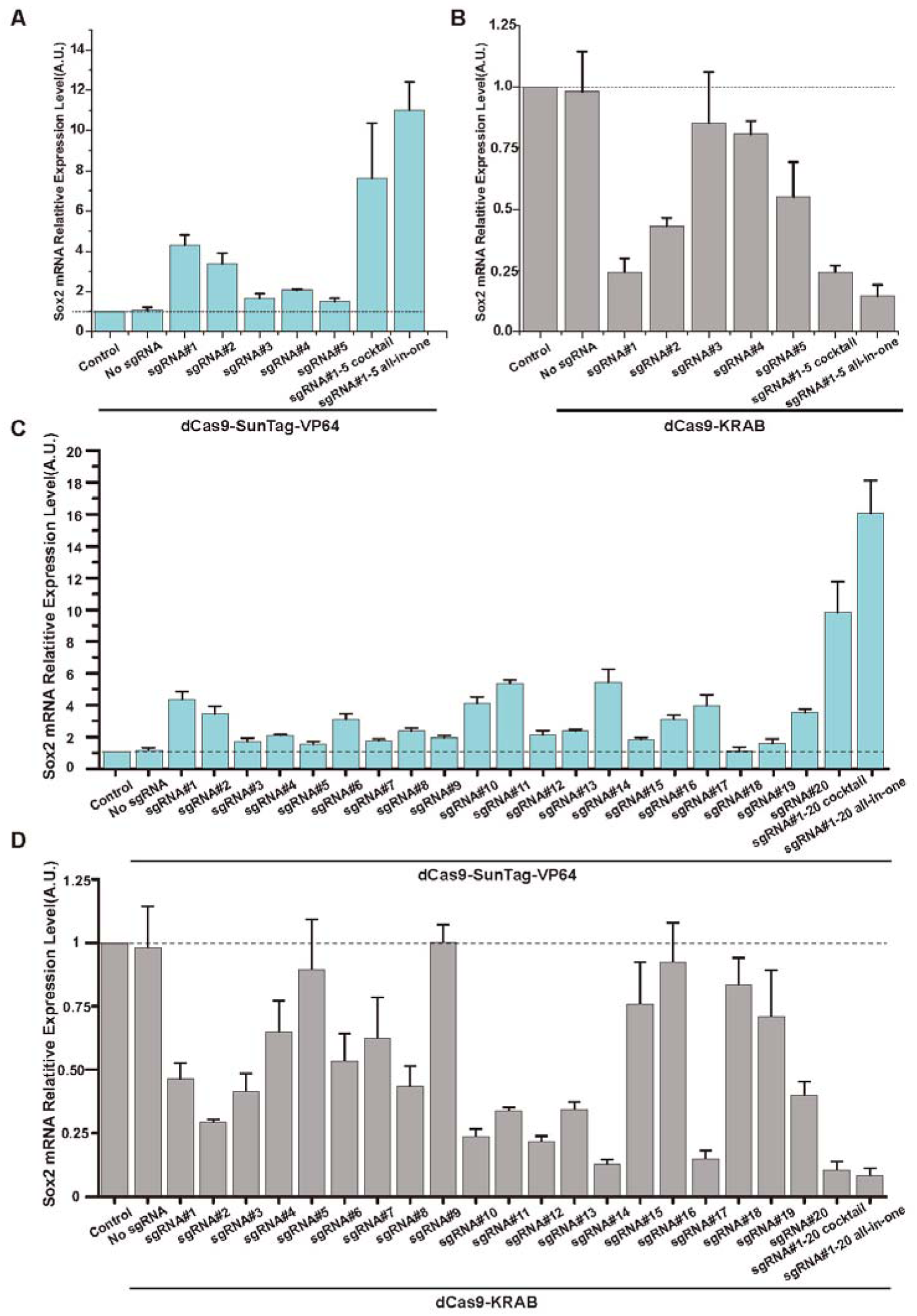
Synergy activation of the endogenous genes by multiplexed sgRNAs targeting one region. (**A**). Synergy gene activation by 5 sgRNAs targeting the promoter region of SOX2 gene. Cells were transfected with either individual plasmid (single or all-in-one) or cocktail of five plasmids. dCas9-SunTag-VP64 are used for the maximum activation efficiency. qRT-PCR were used for the quantification of the activation results. (**B**). Synergy gene repression by 5 sgRNAs targeting the promoter region of SOX2 gene. The same plasmids are used as in (A) except dCas9-KRAB are used instead. Cells without transfection were used as control. (**C**). Synergy gene activation by 20 sgRNAs targeting the promoter region of SOX2 gene. (**D**). Synergy gene repression by 20 sgRNAs targeting the promoter region of SOX2 gene. All values are mean ± s.e.m. with n = 3 biological replicates.

### Coordinated activation and repression of endogenous genes by multiplexed sgRNA

To examine the capability of coordinated gene regulation of our multiplexed sgRNA expression method, we designed 20 sgRNAs each targeting a randomly chosen gene. Cells were transfected with the all-in-one sgRNA expressing plasmid with dCas9-SunTag-VP64 or dCas9-KRAB for activation or repression, respectively. The results indicate that all 20 randomly chosen genes were up-regulated or down-regulated simultaneously (Supplementary Figure 7& Supplementary Figure 8).

At last, we tested the all-in-one plasmid for simultaneous activation and repression of different genes in the same cell. Simultaneous up-regulation of a subset of endogenous genes while down-regulation of another subset of endogenous genes in a defined manner can open up novel opportunities for systems biology as it raises the possibility of manipulating transcriptional networks. We used functional modified sgRNA for this application as different sgRNAs with different aptamers insertion can recruit different transcriptional regulators (Figure 5A). The sgRNAs targeting the first group of 10 genes were inserted with PP7 at their tetraloop and loop2, while sgRNAs targeting the second group of 10 genes were inserted with MS2 at the same position (6). With cotransfection of tdPCP-VPR (25), tdMCP-KRAB, dCas9, and the all-in-one sgRNA expressing plasmid, regulation of different genes was as expected and the regulation result was orthogonal for the corresponding genes (Figure 5B). These results indicate that the multiplexed sgRNA expression strategy can be expanded with modified sgRNA for multi-function applications.

**Figure 5.**
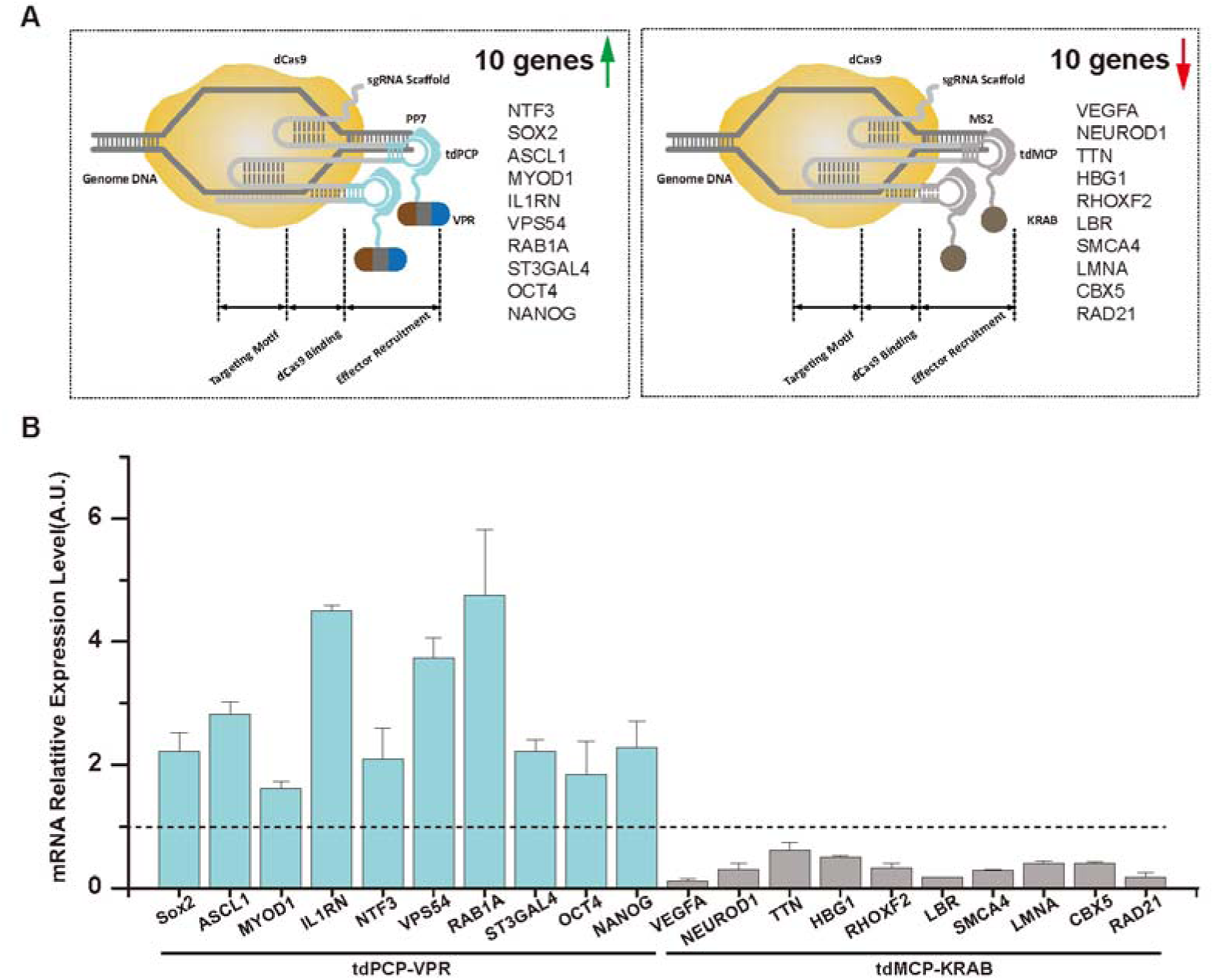
Multiplexed modified sgRNA expression activate and repress multiple endogenous genes simultaneously. (**A**). 10 sgRNA-MS2 (2.02) and 10 sgRNA-PP7 (2.02) were assembled into one plasmid. dCas9, tdPCP-VPR, tdMCP-KRAB and all-in-one sgRNA expression plasmid were cotransfected to cells. 10 genes were activated and 10 other genes were repressed simultaneously in one cell. (**B**). Cells were analyzed by qRT-PCR 2 days after transfection. The genes that are targeted with sgRNA-PP7 (2.02) were upregulated while the gene that are targeted with 10 sgRNA-MS2 (2.02) were downregulated without crosstalk. All values are mean ± s.e.m. with n = 3 biological replicates.

## DISCUSSION

CRISPR based technologies have revolutionized the fields of genome editing and gene regulation. Simultaneous expression and coordinated functioning of multiple sgRNAs will add a new dimension to the evolving CRISPR system, further facilitating our understanding of the physiological and pathological conditions.

Here we have reported a hierarchical assembly strategy for rapid assembly of multiple sgRNA expression cassettes into a single vector. Compared to the current methods, this system ensures high speed, efficiency, capacity and flexibility. Based on Golden Gate cloning, our hierarchical assembly strategy only takes 4-5 days to assemble 20 sgRNA expression cassettes into one plasmid with high fidelity.

Moreover, the hierarchical assembly strategy also offers the flexibility in tuning the number of sgRNAs assembled simply by choosing different combination of amplification primers. Meanwhile, the expression level of different sgRNAs in the all-in-one vector could also be differentiated by using different promoters (19). Additionally, besides the SP CRISPR system, the multiplexed sgRNA expression system can also be used in expressing sgRNAs for other orthogonal CRISPR systems, including SA Cas9 (26), Cpf1 (27) and C2c2 (28). Lastly, a single all-in-one vector can be used for multiplexing operations on the same genomic region, such as imaging, transactivation, repression and chromatin epigenetic modification (29).

In this work, we have demonstrated assembly of 20 sgRNAs in one single vector. Regarding the capacity of the all-in-one sgRNA expression vector, as the upper limit of the sgRNA number that can be assembled is determined by the carrying capacity of the plasmid, there are several possible ways to enhance the system. Firstly, generating multiple sgRNAs from mRNA (30) or tRNA-flanked sgRNAs (31) can make the construct much shorter because only one promoter is needed for the transcription of multiple sgRNAs. Secondly, since the longest fragment in our all-in-one construct is U6 promoter, using other expressing systems will further increase the capacity of our cloning methods. With the improvement of the DNA assembly methods, it is possible that a library of sgRNAs targeting a long DNA sequence can be assembled into one vector, which will greatly aid on the study of structure and dynamics of whole chromosomes or large chromatin domains in living cells. So far, we have only demonstrated the functionality of our all-in-one multiple sgRNA expression vectors in cultured cells. Due to the excessive length and repetition of the multiple sgRNA expression vector, it would be quite challenging to pack it in virus and apply it in *in vivo* studies.

## ACKNOWLEDGMENTS

We thank Dr. Ronald D. Vale (University of California, San Francisco) for providing SunTag plasmids, Dr. Wensheng Wei (School of life sciences, Peking University) for providing sgRNA expression plasmids, Dr. Feng Zhang (Broad Institute) for providing plasmid px330 (Addgene Plasmid #42230), Dr. Scot Wolfe (University of Massachusetts Medical School) for pHAGE-EF1α-dCas9-KRAB (Addgene Plasmid #50919), Dr. George Church (Harvard University) for SP-dCas9-VPR (Addgene Plasmid #63798) and Dr. Wei Guo (Department of Biology, University of Pennsylvania) for providing the MDA-MB-231 cell line. We also thank Ms Xuefang Zhang from the National Center for Protein Sciences Beijing (Peking University) for her excellent assistance with FACS. This work is supported by grants from the 863 Program SS2015AA020406 and National Science Foundation of China 21573013, 21390412, 31271423 and 31327901, for Y.S.

## AUTHOR CONTRIBUTIONS

S.S. and Y.S. conceived and designed the experiments. S.S. and YA.S. performed the cloning experiments. S.S., L.C. and Y.H. performed the FISH experiments. S.S. and X.F. performed the qPCR experiments. S.S. analyzed all the data. S.S. and Y.S. wrote the manuscript.

## Competing financial interests

The authors declare no competing financial interests.

